## Supplementary Figures for "Four layer multi-omics reveals molecular responses to aneuploidy in *Leishmania*"

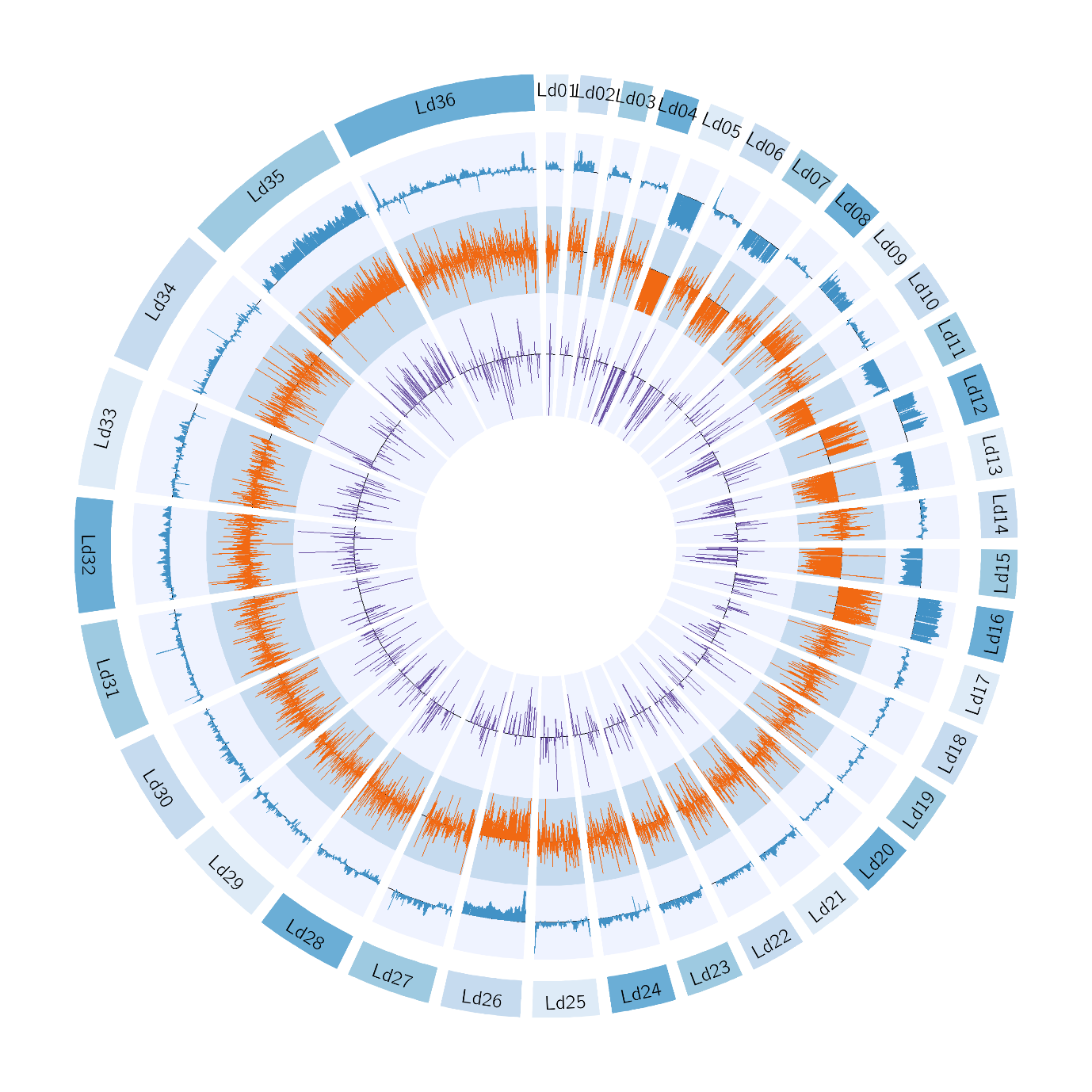

Supplementary Figure S1.A: Circos BPK173 vs BPK288

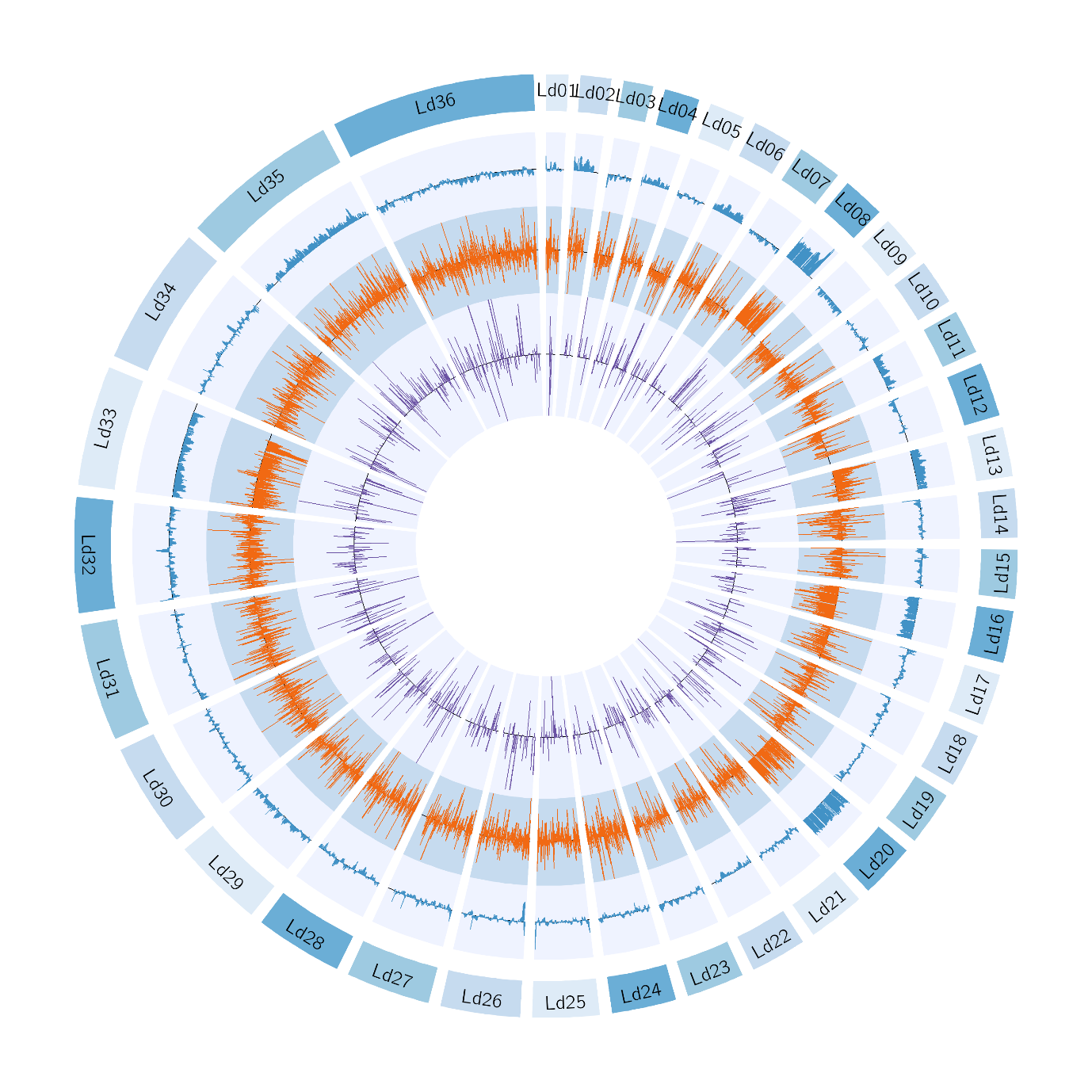

Supplementary Figure S1.B: Circos BHU575 vs BPK282
